# Physics of Self-Assembly and Morpho-Topological Changes of *Klebsiella Pneumoniae* in Desiccating Sessile Droplets

**DOI:** 10.1101/2022.05.04.490658

**Authors:** Abdur Rasheed, Omkar Hegde, Ritika Chaterjee, Srinivas Rao Sampathirao, Dipshikha Chakravortty, Saptarshi Basu

**Affiliations:** Department of Mechanical Engineering, Indian Institute of Science, Bangalore; Department of Microbiology and Cell Biology, Indian Institute of Science, Bangalore

## Abstract

We have investigated the flow and desiccation-driven self-assembly of *Klebsiella Pneumoniae* in the naturally evaporating sessile droplets. *Klebsiella Pneumoniae* exhibits extensive changes in its morphology and forms unique patterns as the droplet dries, revealing hitherto unexplored rich physics governing its survival and infection strategies. Self-assembly of bacteria at the droplet contact line is characterized by order-to-disorder packing transitions with high packing densities and excessive deformations (bacteria deforms nearly twice its original length scales). In contrast, thin-film instability-led hole formation at the center of the droplet engenders spatial packing of bacteria analogous to honeycomb weathering. The varying physical forces acting on bacteria based on their respective spatial location inside the droplet cause an assorted magnitude of physical stress. Self-assembly favors the bacteria at the rim of the droplet, leading to enhanced viability and pathogenesis on the famously known “coffee ring” of the droplet compared to the bacteria present at the center of the droplet residue. Mechanistic insights gained via our study can have far-reaching implications for bacterial infection through droplets, e.g., through open wounds.

## Main

Bacteria orchestrate robust collective responses in dynamic fluid environments by using different physical mechanisms (e.g., suppression of its transport due to fluid shear^1^, oscillatory rheotaxis^2^, swimming against the flow in the wake of curved surfaces/pillars leading to their attachment on the surfaces^3^) that have been developed by evolution over eons. Several recent studies have consolidated the notional importance of the interplay of physical forces in affecting the transport properties of bacteria, their colonization, and survival strategies in novel habitats^4–8^. Unlocking how rugged environments leads to an increase in fitness of the microbes may reveal the physical interactions that cause Anti-Microbial Resistance (AMR)^9^. Opportunistic bacterial pathogens like *Klebsiella Pneumoniae* (KP) are known for their high frequency and diversity of AMR genes^10–12^ and hence are chosen as a model organism in this study. Besides, bacteria spread through complex transmission routes, for example, through aerosols^13^ and fomites^14^, and are subjected to various physical stresses in wild environments.

Bacteria inside droplets experience several physical stresses^15^, such as shear stress due to fluid motion in the environment, starvation stress (caused due to lack of nutrients), desiccation stress (caused due to depletion of water from bacterial cell wall due to evaporation), and stress due to harsh environmental conditions (fluctuating humidity and temperatures), to name a few. The bacteria-laden droplets in contact with any surface will eventually evaporate and lead to a drastic loss in viability of the bacteria^16^ due to desiccation at the end stage of evaporation. However, despite fatal death due to desiccation, few bacteria continue to survive over weeks on inanimate dry surfaces^17^. Studies show that the final survival of bacteria or even virus in an evaporating droplet depends on its ability to withstand drying-induced stress^18–20^. The bacteria on the dried residue of droplets also form patterns, including “coffee rings”; however, motile bacteria can also alter the deposition patterns by evenly distributing the bacteria on the contact surface of the dried residue of the droplet^21^. The wettability of the bacterial surface/cell wall also plays a vital role in forming deposition patterns and the adhesion characteristics on the surface^22,23^. Order to disorder transition in the coffee ring deposit is observed in both spherical and rod-shaped inert particles^24,25^. The disordered region is characterized by a random packing pattern and dispersion in the Voronoi area^26^. However, it is unclear if an active matter like bacterial cells would exhibit a pattern and trend in packing at the coffee ring.

There are significantly fewer works on the effect of desiccation on the survivability of bacteria^27–31^. Even these works have only studied what happens to the overall viability without considering the fate of bacteria (i.e., its morpho-topological changes, the viability variation within different regions of the deposit, the effect of self-assembly). In the present work, we will explore how the physical mechanism of self-assembly of KP in evaporating droplets favors a few bacteria to survive the adverse condition. Investigating the complex patterns formed by the evaporating droplets containing bacteria that may enhance the survivability and pathogenesis of bacteria, in the present article, we study the physics pertaining to the self-assembly-based survivability mechanism of KP in evaporating sessile droplets.

### A. Bacterial self-assembly in evaporating droplets

Particle aggregation towards the contact line on pinned sessile droplets evaporating in an ambient environment is a well-known phenomenon^32^. The colloids suspended in the droplet are driven along with the flow created to replenish the evaporating water at the edge of the water droplet, resulting in a coffee ring-type deposition. In contrast, bacterial aggregation is much more complex than inert particles. For example, bacteria are soft/deformable, have semi-permeable cell walls, interact, form colonies/combined structures, etc. In addition, droplets containing bacterial suspension exhibit pinned contact line for the entire evaporation lifetime on hydrophilic substrates^33–35^.

In our experimental set-up, we placed a 1 μl droplet of milliQ, suspended with the KP bacteria, on the clean glass surface, as shown in Fig. 1 (c) (refer to the “Droplet Evaporation” in the Methods Section for the detailed protocol). As observed from the live-cell imaging, most of the KP bacteria adsorb at the surface of the droplet (as represented in Fig. 2 (a)) due to the high electric charge on the bacterial surface (refer to Table 1 in the supplementary information). Several bacteria faithfully follow the flow inside the droplet, laterally stacking themselves at the droplet rim (see Video 1, Fig 2. (b)).

**Figure 1.**
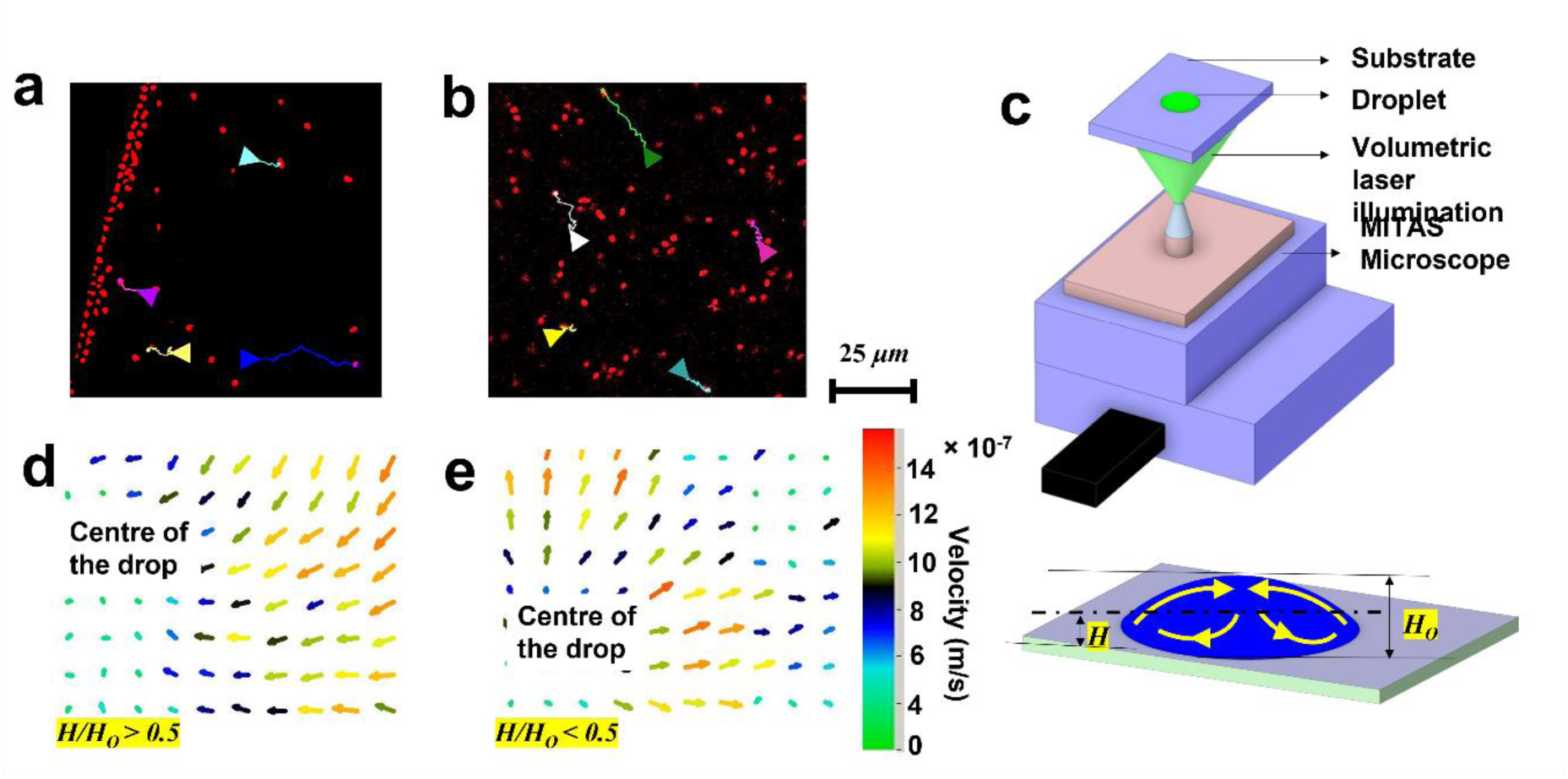
Live Cell tracking of bacteria and hydrodynamics inside the droplet laden with *KP*. Snapshot of live cell image of the KP suspension droplet (at *t/t*_*f*_∼ 0.1) at the (a) edge of the droplet, (b) center of the droplet. The colored arrows in (a) and (b) show the direction of transport of KP, and the corresponding colored line is the path line followed by the respective bacteria for 10 seconds. (c) Schematic representation of the μ-PIV experimental set-up. Vectors of velocity (d) at *H/H*_*0*_ ∼ 0.1 (e) at *H/H*_*0*_ ∼ 0.6. The scale bar for the velocity vectors for (d) and (e) is adjacent to (e).

**Figure 2.**
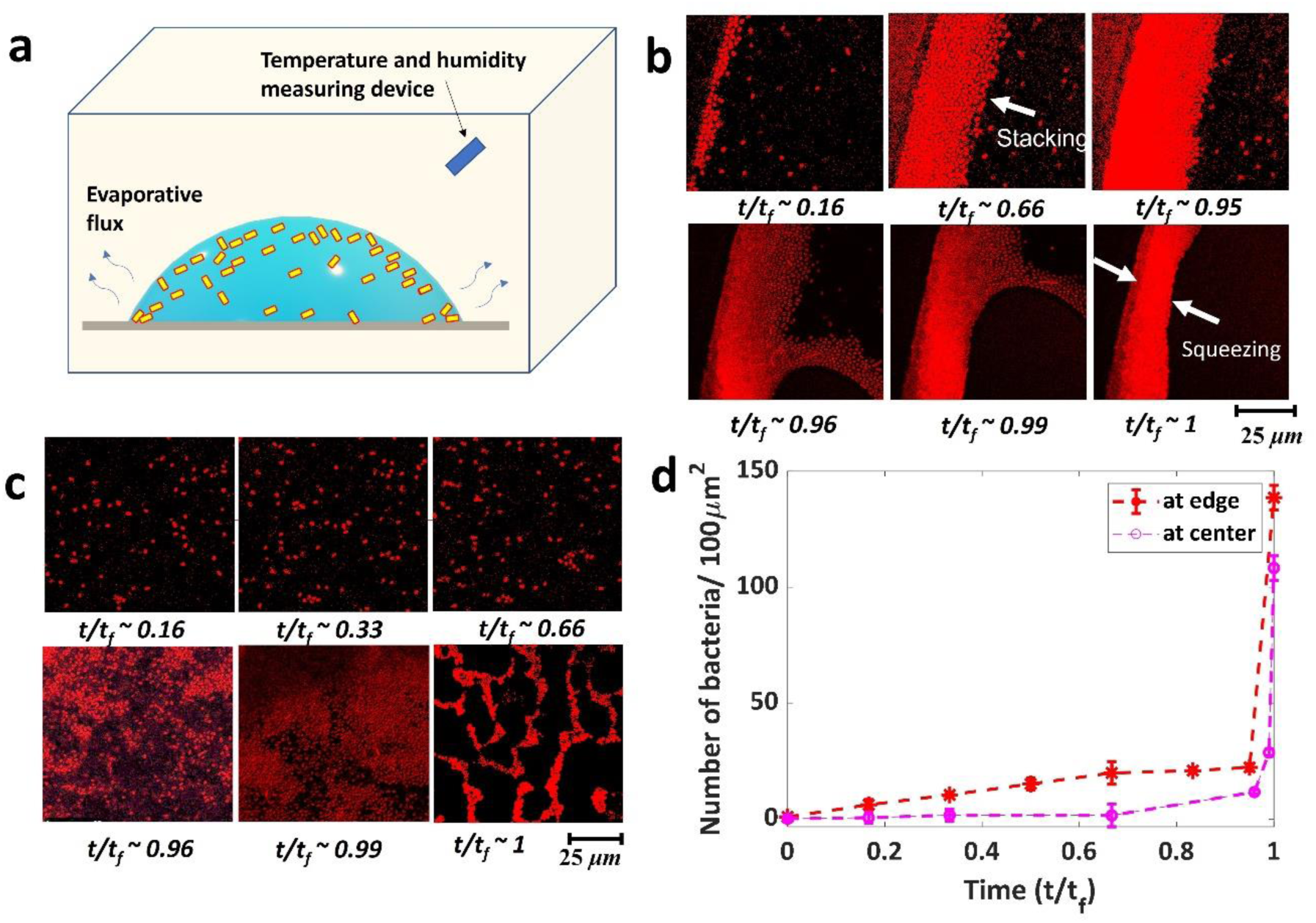
Spatio-temporal dynamics of self-assembly of *Klebsiella Pneumoniae* (KP) in evaporating droplets. (a) Schematic of an evaporating droplet containing the KP bacteria in controlled atmospheric conditions. Temporal sequential live-cell images of KP deposition near the glass surface (at a plane 10 μm from the surface), (b) at the edge, (c) at the center of the droplet. (d) Bacterial count scaled/100 μm^2^. For time *t/t*_*f*_=1, the bacterial number is counted from the SEM images (as the individual bacteria is not distinguishable in the fluorescence images).

Although evaporation-driven flow (EDF) inside the droplet exists, all bacteria may often not faithfully follow the flow^36^. Live-cell imaging reveals the real-time transport of bacteria in the droplet near the glass surface (at a 10 μm distance from the glass surface); Fig. 2 (a) and (b) show the path line followed by the individual bacteria. Several bacteria move towards the edge of the droplet; however, few move away from the edge (refer to Fig.1 (a) and Video 1). Few bacteria in the central region get adhered to the surface and wiggle there, while few others move in an arbitrary direction. Further, we study the flow inside the droplet to understand its role in the transport of KP in the droplet. The Micro-Particle Image (μ-PIV) Velocimetry technique is used to quantify the flow inside the droplet, and the schematic representation of the experimental set-up is shown in Fig.1 (c) (refer to the Methods section for the details of the μ-PIV experiment).

Our investigations show circulatory flow in the bacteria-laden Milli-Q water droplet. The flow is radially inwards (i.e., towards the center of the drop), as shown in Fig.1 (d). Near the glass surface, the flow is directed radially outwards (see Fig.1 (e)). Since the KP bacteria used in the experiments are non-motile, the bacteria moving away from the edge of the droplet (in Fig.1 (a)) are, in reality, moving upwards along with the recirculatory flow inside the droplet. Near the end of evaporation, the bacteria that has not settled in the coffee ring gets deposited in the rim of the local dewetting hole front, as discussed in section C.

As shown in Fig. 2 (b), the ordered packing of KP is similar to the ordering of rod-shaped particles previously observed by Dugyala et al.^37^. As the height of the droplet becomes thinner, the evaporative flux at the contact line increases. Subsequently, the flow velocity inside the droplet towards the contact line also increases. Studies have shown a rapid increase in velocity near the end of evaporation, usually referred to as ‘the rush hour effect.’ The resulting deposition at the coffee ring generally exhibits an order to disorder transition^37–39^. We observe a similar order to disorder transition self-assembly in the case of KP bacteria.

The bacterial number density in a region is calculated by considering a region of interest in the image (for a given instant) and subsequently counting the number of bacteria in the given region. The values plotted in Fig.2 (d) are the averages of bacterial count in six arbitrary regions at any given instants, and the error bar corresponds to the standard deviations of the same. This is scaled/100 μm^2^ in order to have a substantial number of bacteria for a given region. However, in the end stage of evaporation (*t/t*_*f*_= 1, where *t* is the image at any given time instant and *t*_*f*_ is the total time of evaporation of the droplet), the bacteria is squeezed and very closely packed such that it is not possible to obtain the bacterial count in the given region from the live-cell image. Hence, the bacterial count at *t/t*_*f*_= 1 is calculated from the Scanning Electron Microscopy (SEM) images (See Fig.S1 in the supplementary information). It is observed that the bacterial number density in a given region at the droplet rim increases drastically with time, leading to very close packing of the bacteria (refer to Fig. 2 (b) and (d)). At the center of the drop, we see a sudden increase in the visible number of bacteria during the end stages of evaporation. The focus is adjusted to visualize a section plane of a droplet near the substrate (at a plane 10 μm from the glass surface). Since many bacteria reside near the air-water interface of the drop, as the drop height reduces, the bacteria move along with the open surface of the droplet towards the substrate and thus become visible in the live-cell imaging in the end stage of evaporation.

The thin film of liquid and bacteria residue at the end stage of the evaporation of the droplet (*t/t*_*f*_∼0.99) undergoes instability; subsequently, capillary forces dominate, leading to the formation of holes and compact packing of KP as shown in Fig.2 (c). The bacterial number density at the center of the droplet residue (for *t/t*_*f*_=1) is calculated from the SEM images in the region where the bacteria aggregates disregarding the region of holes (see Fig.2 (c)). Fig. 2(d) shows that the compaction of KP happens in the last one percent of the total evaporation times, and the self-assembly is dominantly the result of capillary forces and thin-film instabilities at the end stage of evaporation. This phenomenon in the case of KP is explained in sections C and D.

### B. Ordered to dis-ordered transitions and simultaneous squeezing and buckling of bacteria on “coffee ring”

Based on the type of self-assembly, we have classified the regions within the droplet residue into three types: 1) ordered region near the edge of the droplet residue – referred to as “Outer Edge” as shown in Fig.3 (a) and (b). 2) the disordered region – referred to as the “Inner edge” as shown in Fig.3 (a) and (b). 3) the central region of residue – is referred to as “honeycomb weathering” Fig.5(a). An aerial view of the bacterial deposits captured from SEM has been used to analyze the size and bacterial ordering. The bacteria profile is manually marked using a digital pen to precisely detect the bacterial edges while processing the image (Fig.3 (g)). ImageJ software is used to binarise the image, and the ‘analyze particle’ plugin is used to get the size and position characteristics.

We define the maximum and minimum ferret length ratio as the aspect ratio (ratio of length (L) to the width of the bacteria (W)) of the bacteria. Aspect ratio and vertical height are higher for the bacteria that got squeezed laterally. The Aspect ratio of bacteria is averaged radially from the edge (see Fig.3 (b)). The radial distance from the edge, R, is normalized with the thickness of edge deposition (the coffee ring thickness), R_e_. Three different deposits have been used to get the statistical mean of the plot shown in Fig 3 (c). The deposits from separately grown cultures have also been tested to ensure repeatability. The aspect ratio is higher at the outermost edge than in the central region for all the cases. The aspect ratio is highest at R/R_e_ between 0.35 to 0.5. Bacteria in this region are highly jammed between the bacteria that settled at the outermost edge and the incoming bacteria during the rush hour and squeezing effect, as discussed earlier. The bacteria in the inner edge are in a wide range of aspect ratios depending on the configuration of the bacteria. Bacteria at the inner edge may be folded due to excessive desiccation stress and attain U shape^40^.

**Figure 3.**
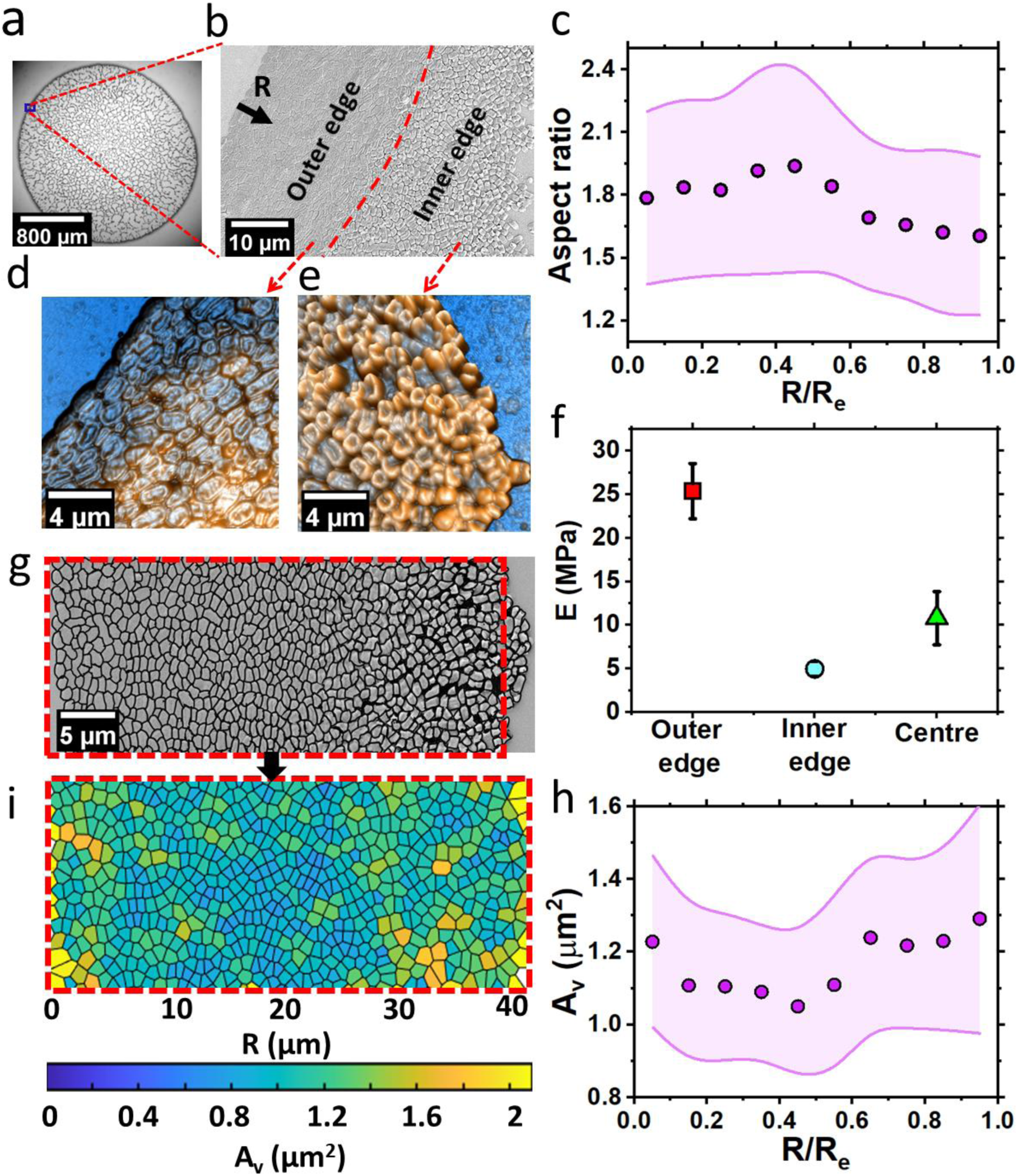
Bacterial Self-Assembly at the edge of the droplet. (a) Optical image of the dried residue of the KP droplet. (b) SEM micrograph of the KP deposition at the edge/rim of the deposit. (c) The plot of the radially averaged aspect ratio of the bacteria as seen from the top view vs. the normalized distance (R/R_e_) from the outer edge. Where R is the radial distance from the edge of the contact line, and R_e_ is the thickness of edge deposition. Atomic force microscopy images at the (d) outer edge and the (e) inner edge of the precipitate as shown in (b). (f) The plot of the Modulus of Elasticity values (E) for bacteria at the outer edge, inner edge, and center. g) SEM micrograph on the edge of the deposit, bacteria marked with black border for centroidal Voronoi tessellation processing. (i) Voronoi cells of the corresponding bacterium shown in. g) color marked according to their size. (h) The Voronoi area of the bacterial deposits was averaged and plotted with normalized distance from the edge R/R_e_.

Atomic force microscopy, AFM image (Fig. 3 (d) & (e)) confirms this observation and gives a clear picture of the bacterial morphology at the inner and the outer edge. The bacteria at the outer edge are packed with their major axis along the periphery, whereas the bacteria at the inner edge exhibit a random arrangement. Marin et.al^24^. observed order to disorder transition in inert sphere particles, and Dugyala^37^et.al^25^. observed this in colloidal rods. The thickness of the bacterial deposits at the edge varies from 300 nm at the outermost edge up to 700nm in the highly squeezed region. At the same time, the folded bacteria at the inner edge showed a height in the range of 350 to 500 nm. Height variation in the outer edge can occur because of variation in squeezing and water content in bacteria due to variation in the level of desiccation. Whereas in the inner edge, this variation is due to the degree of folding up of the bacteria, the bacteria that have folded up more show higher height. Folding of bacteria is a preventive measure to avoid desiccation. The bacteria try to reduce the exposed area for desiccation by attaining a coccoidal shape ^40^. The bacteria at the inner edge folds because the lateral surfaces are also exposed for desiccation, unlike the bacteria at the outer edge region enclosed by the closely packed adjacent bacteria hindering evaporation from its lateral sides. The bacteria at the outer edge are closely packed, so they are constrained to fold up like those in the disordered region. Folding is neither seen in the outermost first line of the bacteria on which one of the lateral surfaces is exposed to the ambient. We hypothesize that the bacterial cell-substrate bonding might be higher here due to prolonged cell-substrate interaction, which would have prevented cell folding in the lateral direction.

The force-distance spectroscopy analysis done on the bacterial samples through nano indentation (refer to methods section for details) at various regions in the deposit reveals the elasticity of the cell membrane of the bacteria. The Modulus of Elasticity (E) of the bacterial membrane is obtained using the Oliver Pharr model. The results show that the outer edge bacteria are stiffer than the other bacteria. The squeezing effect on these bacteria leading to very close packing, has increased the stiffness (see Fig. 3 (f)) by nearly one order. E of the bacteria is 25 ± 3 MPa, 5± 1 MPa, and 11 ±3 MPa at the outer edge, inner edge, and central region, respectively. The variation in the elasticity modulus can be attributed to the level of desiccation and ordering dynamics induced gap between the bacteria based on the location.

In addition, the difference in the cross-sectional area normal to the indentation direction would also be a factor as the bacteria is twice as stiffer in the circumferential direction than in the axial direction.^41^ Deng et al. showed that axial Modulus E = 23 ±8 MPa, is less than half the circumferential Modulus E = 49 ±20 MPa. In the current study, we obtained the elasticity modulus of the exposed top surface by AFM cantilever tip indentation for all cases. In the case of outer bacteria lying with their axial direction along the periphery of the droplet deposit, the top surface indentation gives the circumferential Modulus. The bacterial cross-section in the axial direction is lesser than in the circumferential direction, and such is the elasticity modulus. The inner edge bacterial cross-section gets reduced by folding up, which decreases elasticity modulus. Correspondingly, we hypothesize, that the bacteria with less water content is less stiff and deforms more at a given force during indentation. The reasoning is further supported by the maximum unbinding force (force required to detach the AFM probe from the sample during retraction) on the indenting probe shown in Fig.S2. The bacteria deposited at the outer edge show the highest unbinding force (adhesion force) compared to the bacteria at the other sites. Bacterial cells secrete protein across their membranes upon contact with other surfaces^42,43^. The variation in adhesion force on the bacterial surface can be attributed to the secretions from bacterial cells upon interaction with the substrate. Bacterial cells at the outer edge are in close proximity with the substrate for a longer duration before desiccation, whereas the bacterium at the central regions gets deposited much later. Shear stress imposed on the bacteria in different regions could also be an essential factor in expressing adhesive substances through its membrane which inturn reduces desiccation.

The centroidal positions of each bacterium obtained through image analysis are used to study the bacterial ordering. The bacterial ordering is analyzed by centroidal Voronoi tessellation using MATLAB. The Voronoi cells in Fig. 3 (i) show the Voronoi area (A_v_) varying from 0.5 μm^2^ to 1.85 μm^2^. The Voronoi area of the cells is obtained by processing the SEM image Fig.3(g), where the bacterial profile is marked manually for discrete identification of cells. The average Voronoi area is 1.1 μm^2^ at the outer edge and 1.21 μm^2^ at the inner edge. The Voronoi area reduces inwards in the outer edge region where the bacterium is more squeezed. In comparison, the Voronoi area increases towards the disordered region. For inert non-deformable particles, it is observed that the Voronoi area shows a dispersion (sudden increase in the Voronoi area) from the order to the disordered region^24^. Evidently, such dispersion is not observed in the bacterial deposition. There could be three causes for this: the squeezing effect during the hole growth, the capillary force leading to packing during the evaporation of the mesoscopic thin layer, and the folding of the bacteria.

The supplementary Figure (Fig.S2) shows the overall picture of bacterial deposition at the edge. The deposit thickness increases from the outer edge and reaches a maximum at R/R_e_ of around 0.4 to 0.5. There is a strong relation between the increase in thickness, the decrease in the Voronoi area, and the increase in aspect ratio. The squeezing effect leads to morphological changes in the bacterial cells resulting in the increment in height and reduction in the area of contact with the substrate to conserve the volume. Moreover, due to lesser exposed surface area and desiccation, these bacteria would have more water content and have higher volume than the edge bacteria and the ones with a gap between them (inner edge). The height variation of bacterial deposits in the central regions remains less the same, around 200-300 *nm*(Fig.S3) at a different location depending on the local ordering of bacteria.

### C. Thin-film instability leading to honeycomb weathering

Not all bacteria in the droplet get settled at the ring. Depending on the concentration, interface, and evaporation dynamics, a significant proportion of bacteria would remain dispersed in the liquid film. In the current study about 50 to 60 % of the bacteria gets deposited on the edge and the thickness varies from 38 to 44 nm. Liquid thin-film dynamics determine the final deposition of bacteria not settled in the coffee ring. The perturbations in the liquid thin film would cause rupture, hole growth, and resulting island formation^44^. The size and shape of these islands depend on the liquid properties and size, shape, and interfacial properties of the colloids. The bacteria come closer, leaving a region of low liquid film thickness where the rupture would occur. ^45–47^.

The deposition of bacteria in the central regions follows a sequence of events discussed in this section. At shallow film thickness, the contact line de-pins, leaving a fine mesoscopic layer of thin liquid film containing bacteria distributed throughout. The depinning is followed by the thin film instability leading to the cellular pattern formation. Due to thin-film capillary instability, the liquid film thickness varies along the wetted area. These perturbations get enhanced due to the presence of bacteria. Fringe patterns were observed as shown in Fig 4 a) through the reflection interference microscopy at low film thickness. Figure 4 shows the thin-film variation of the bacteria-laden droplet at various time instants and the phenomena of the dewetting process of the liquid layer at the conclusive phase of evaporation. Here we take t_start_ as the time when depinning has started and t_i_ is the instantaneous time which is t_start_ when depining has initiated. At the initial time instants (t_i_ = t_start_ +3.28s), densely spaced fringe patterns were formed in the form of dark and bright bands, as seen in Fig.4 (a). These patterns signify the presence of a thicker liquid layer, and from the initial observance of the fringes, the contact angle of the liquid layer is estimated at approximately ≈5.6°. As time progresses, the fringe width evolves to wider patterns, and instigation of circular patterns of the liquid layer is evident nearer to the contact line of the droplet. The formation of multiple circular patterns in the liquid layer for (t_i_ *≈* t_start_ + *29*.*20)* represents the thinning of the liquid layer compared to the initial time instants that result in a reduction in thickness of the thin film. The observance of multiple circular rolls is slightly distant from the leading edge. This phenomenon is predominant due to the deposition of more bacteria along the droplet’s edge.

**Figure 4.**
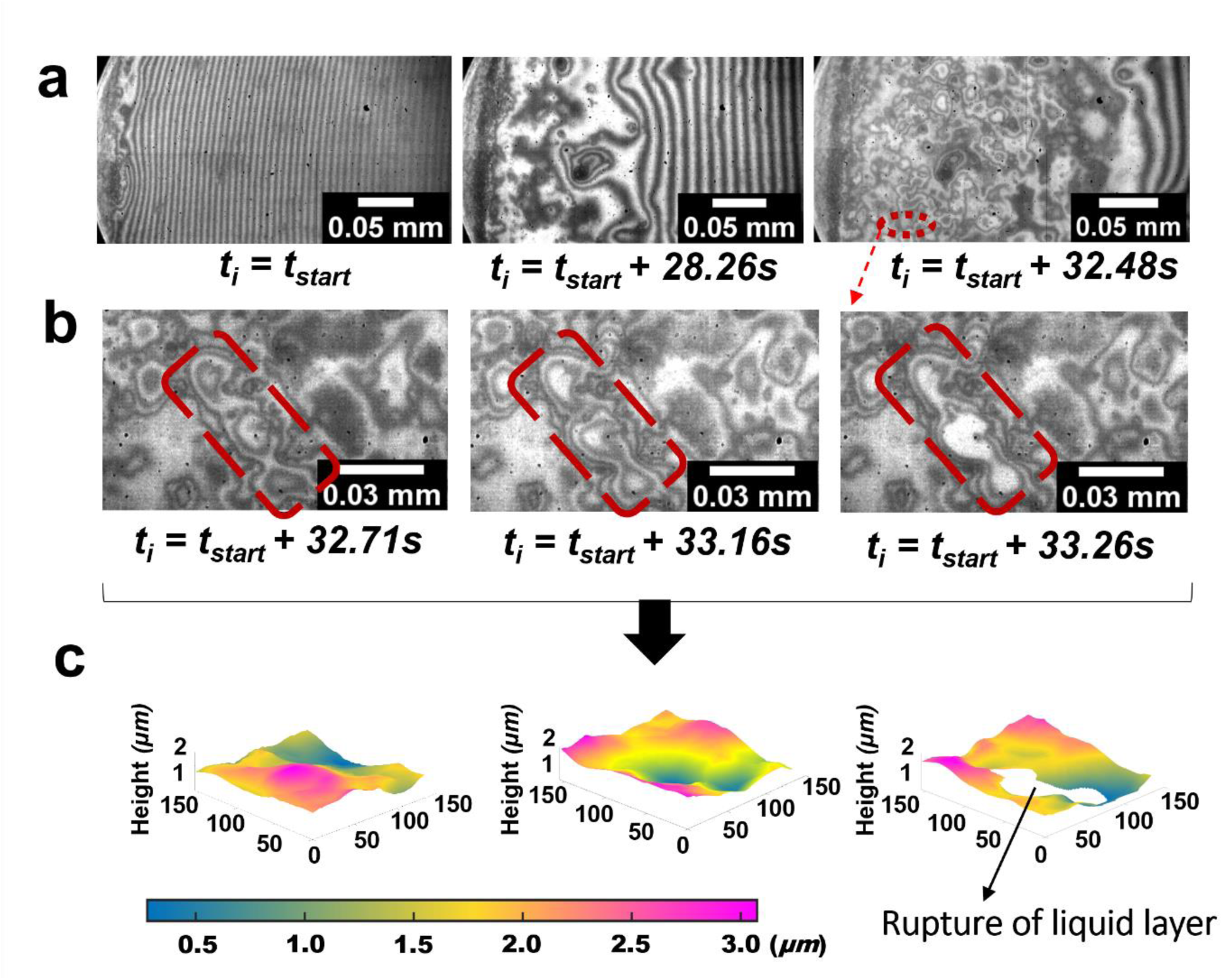
Thin film instability at the center of the droplet. (a) The liquid layer at varying time instants during evaporation of the droplet (b) Insights of the de-wetting process in the thin-film layer (c) Variation of the thickness profiles along with the de-wetting phenomena.

The pinning of droplet without receding the liquid layer consents to the existence of the thin film. Once the thickness of the thin film is reduced, the evaporation of the liquid layer surrounding the bacteria elucidates the variation in thickness resulting in the formation of multiple roll structures and the onset of the dewetting process of the liquid layer. The rupture of the thin film and the sequence of variation in thickness of the liquid layer at the selected regions of the initial de-wetting process are presented in Figs 4 (b) and (c). The formation of multiple rolls at a specific time of 29.43s and adjoining of the rolls in the course time till the rupture of the thin film is evident from Fig. 4 (b). The rupture of the liquid layer is observed from the interferometric images at t_i_ = t_start_ +29.98s. The associated two-dimensional distribution of the height profiles and variation of the thin film was presented in Fig. 4 (c). During the t = 29.43s, the presence of multiple rolls (in the selected yellow regions) shows the increase in the height of the thickness profile, nearly having the maximum height of 3 µm.

In contrast, the adjoining regions of the rolls represent the reduction in thickness variation nearly to 1µm. Similarly, at t_i_ = t_start_ +29.98s, the distribution of the height profile surrounding the rupture region of the liquid layer varied from 0.5 to 1.5 µm as the liquid layer recedes to the adjacent regions. Therefore, the formation of multiple roll-structures (as in t_i_ = t_start_ +29.20) leads to multiple de-wetting locations regions in the fluid layer, leading to multiple streaks during the droplet’s evaporation (deposition of bacteria) from the droplet edge to the center of the droplet. Since the droplet de-pining initiates from the edge, the hole formation starts from the edge. The hole formation progress towards the central region as the film thickness decreases, as shown in Fig. 5 (a). While the hole near the droplet edge keeps growing, another hole forms adjacent to the droplet center. The growing front from two holes merges, creating an island of bacteria.

**Figure 5.**
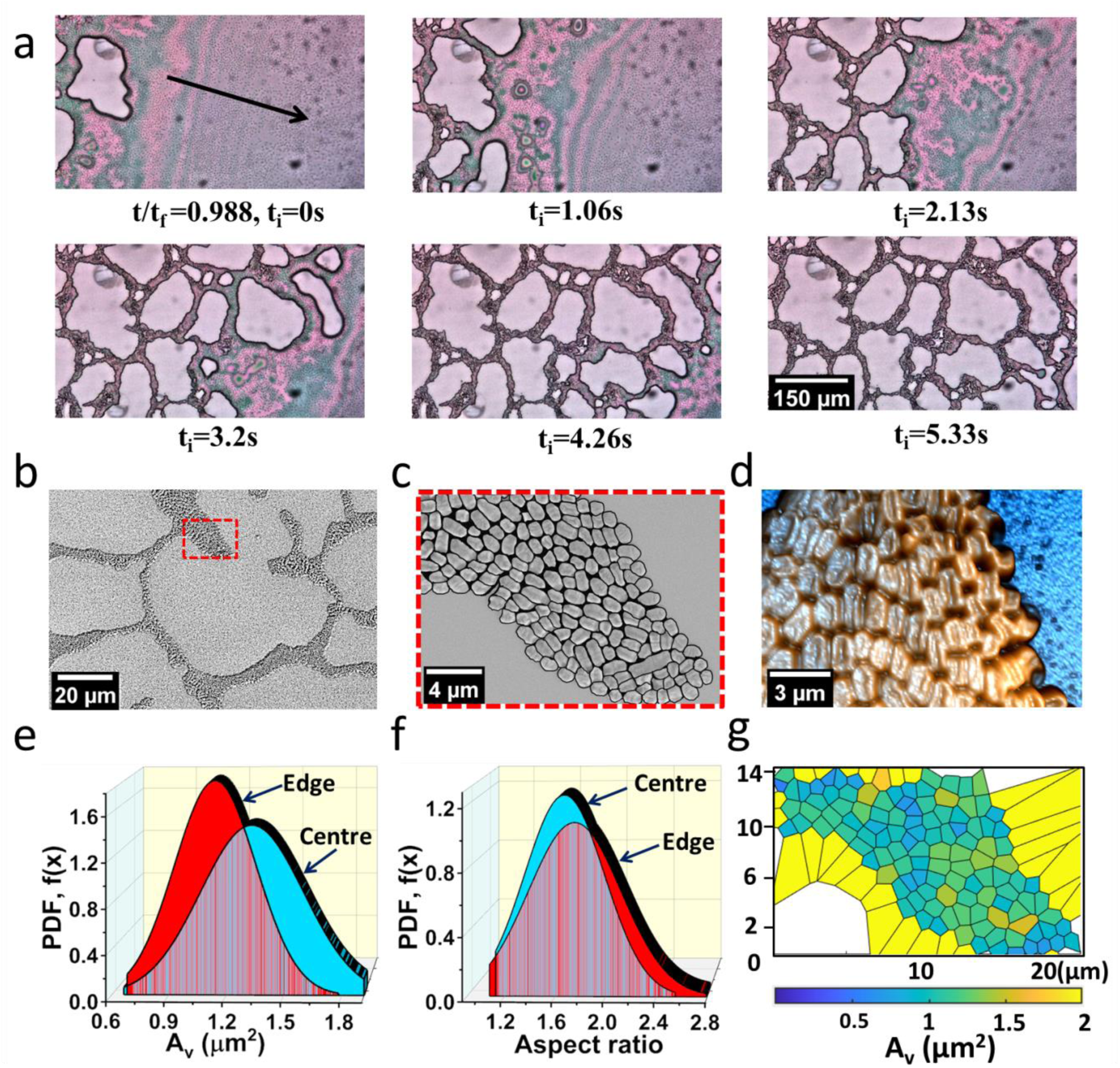
Bacterial Self-Assembly at the central region of the droplet. (a) Spatio-temporal evolution of the liquid film leading to the rupture and hole growth resembling a cellular pattern (b) SEM micrograph of a cellular structure at the central region (c) Zoomed in SEM micrograph with marked bacterial edge profile, represent the bacterial ordering in the cellular structure (d) AFM image shows the deposited bacterial morphology and self-assembly at the central region (e) Plot comparing the Voronoi Area at the central region to that of the edge of the deposit through Probability density function, PDF, f(x) plot (f) plot of PDF of aspect ratio (viewed normal to the substrate) of bacteria at the edge and center (g) Color-marked Voronoi cells of the bacterial deposits corresponding to (c).

**Figure 6.**
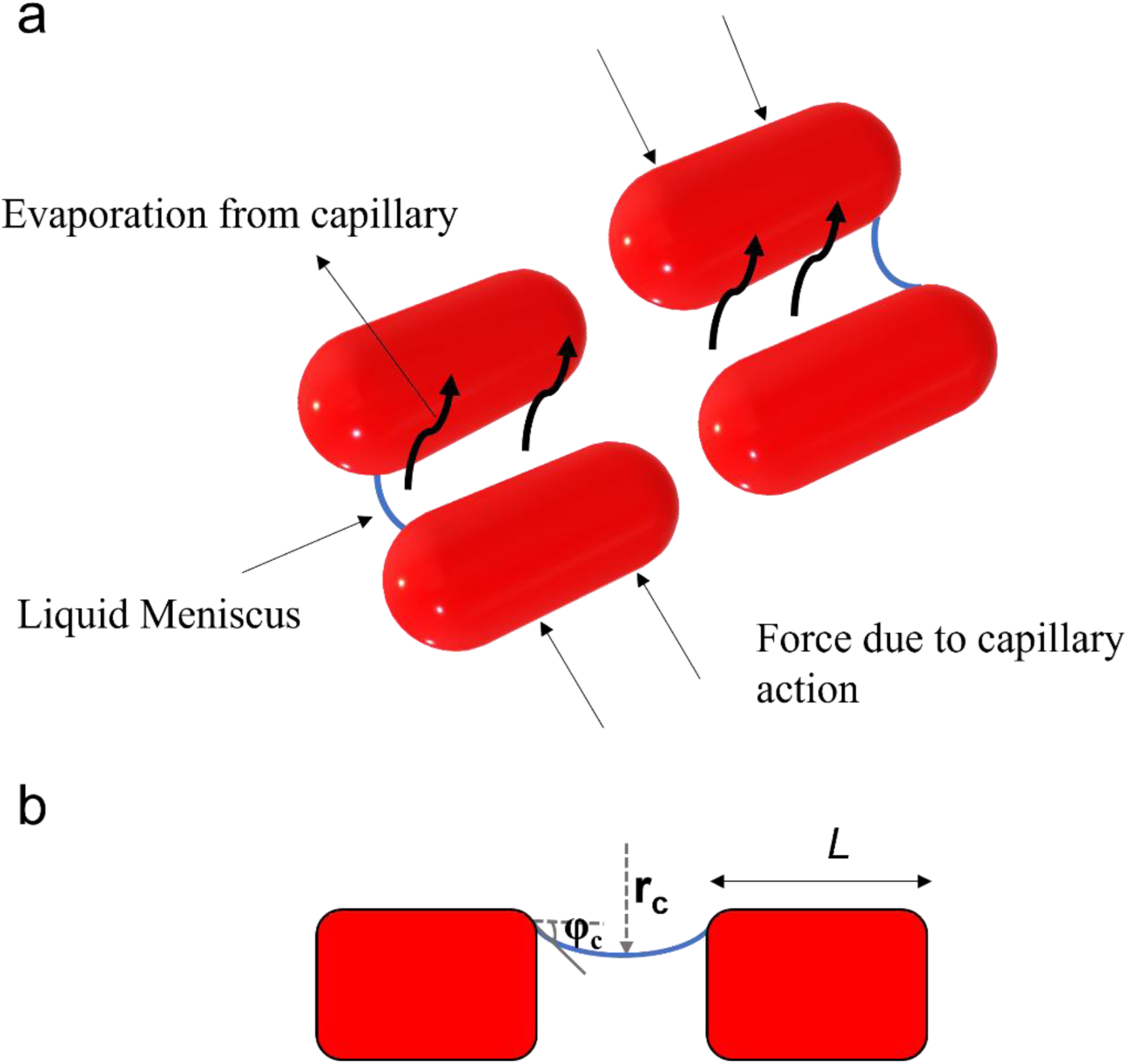
Mechanistic insights on bacterial aggregation. Schematic representation of (a) formation of a meniscus leading to the capillary attraction between the bacteria at the end stage of evaporation, (b) Side view of the liquid meniscus in between the bacteria

**Figure 7.**
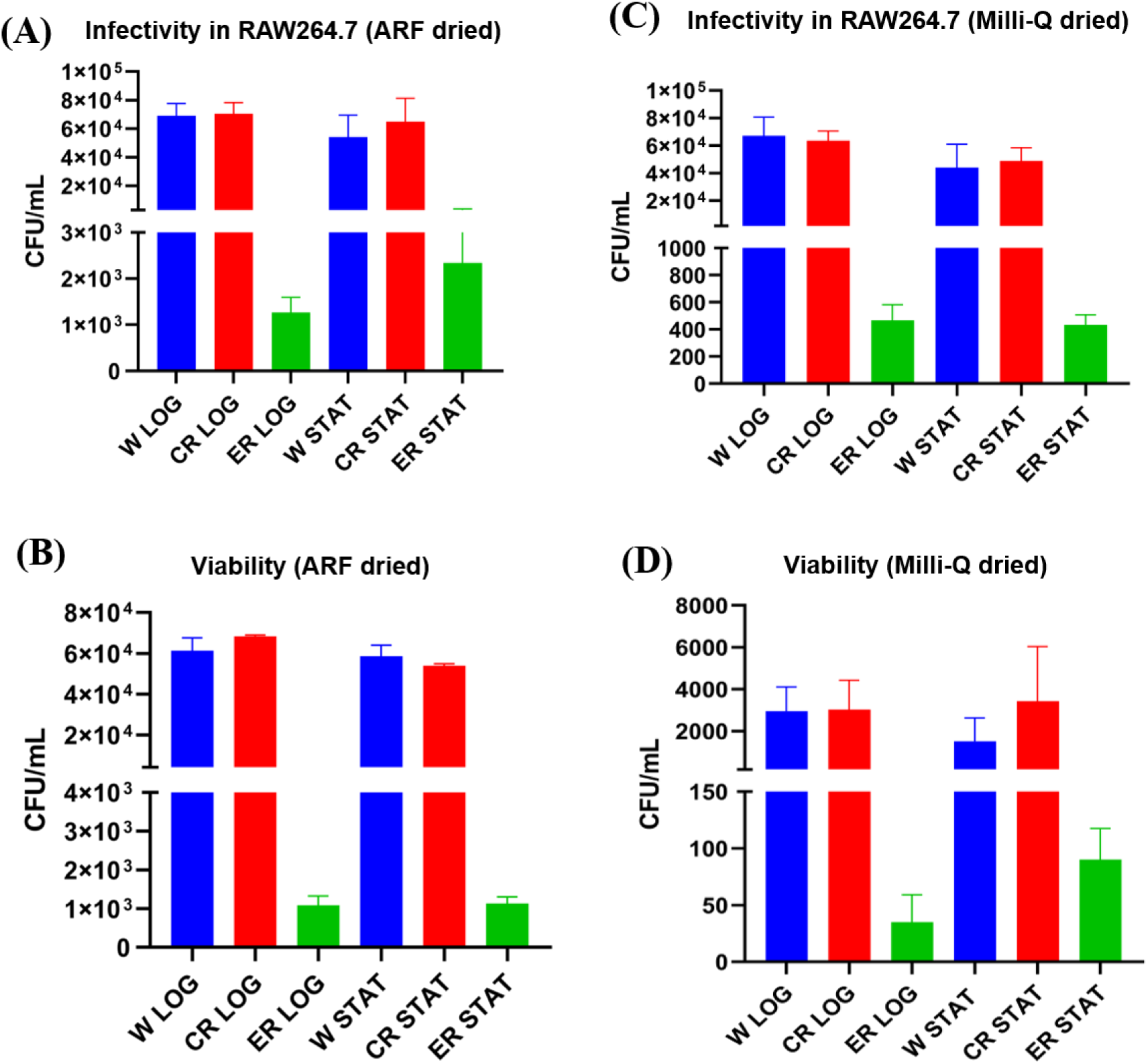
Bacterial survivability and pathogenesis. (A) Infectivity of the bacteria dried in surrogate respiratory fluid (SRF) droplet in murine macrophages RAW264.7 cells. (B) Viability of bacteria dried in SRF droplet. (B) Infectivity of the bacteria dried in Milli-Q water droplet in murine macrophages RAW264.7 cells. (B) Viability of bacteria dried in Milli-Q water droplet. All experiments are repeated independently three times (N=3); error bars in the plot correspond to the standard error of the mean.

Due to perturbations on the thin film, the hole formations will either increase the surface energy or decrease it. For a negative change in energy, the hole continues to grow, and for a positive change in energy, the hole shrinks. Using the energy minimization criterion for a quasi-static system of hole radius r_o_ and inner hole distance D, the change in energy of a perturbed system ΔE can be given as^44^,

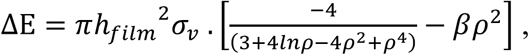

Where,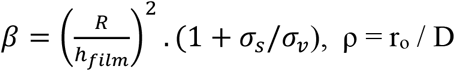

For β<19.5, holes of any size will shrink, and the film will become flat. At β>500, holes will grow for most of the range of ρ. Infinitesimally small holes will shrink at any value of β. *σ*_*v*_ represents the film-vapor surface energy *σ*_*s*_, is the film-substrate surface energy. The major players in determining the energy change are the hole size, inter-hole distance, and film thickness. At high film thickness, smaller holes produce a positive change in energy, resulting in shrinking while larger holes could grow. At a film thickness of order 10 μm, holes of size r_o_>3.2 μm will create a negative change in surface energy and start to grow in the current case.

The instantaneous hole growth at the edge and the central region is compared in Fig.S4 The pinned contact line, and the edge deposits impede the growth of holes near the coffee ring. The bacterial cell assembly at the edge is squeezed by the growth of holes adjacent to it. The cells cannot move further due to the pinning of the contact line. Thus, the hole growth is restricted toward the coffee ring. However, the holes can still grow towards the center of the droplet. The normalized hole area exhibits a linear variation in Fig.S4.

The holes formed at a location further away from the edge can grow with lesser restriction initially. As the adjacent holes start to impinge on each other, growth is slowed down. The normalized hole area variation in the central regions fits very closely to the logarithmic curve (Fig.S3). Here, the growth rate reduces exponentially with time, which shows that the dominant mode in the hole growth mechanism is the surface diffusion^44^. Whereas for the holes near the coffee ring, the growth rate is nearly the same as dictated by the evaporation kinetics^44^. The hole growth rate for an evaporation dominated case is, 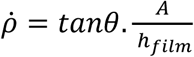 Where θ is the equilibrium wetting angle, A is the gas-phase transport coefficient, defined as, 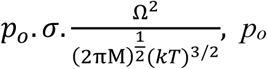 is the equilibrium vapour pressure on the film surface, σ is the surface energy per unit area of the film/vapour interface, Ω is the atomic volume, M is the atomic weight, and kT is the thermal energy. 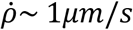 which is of the same order as the growth rate observed in the experiments for the holes. The holes at the central region have an initial velocity two orders higher than that calculated, which drastically reduces to the order of 1*µm*/*s* in less than half the growing time. The total growing time, shape, and size of holes vary randomly. The position of the hole inception, the inter-hole distance, and bacterial density distribution around the hole are the significant factors influencing the final hole size and shape, along with the fluid properties.

Habibi et al.^48^ studied the pattern of particles in dewetting thin films and proposed a simple model for predicting the dewetting velocity,

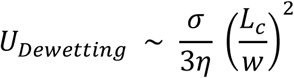

Where σ is the surface tension of the liquid film, η is the effective viscosity of the thin liquid film, w is the characteristic size of the hole, and *L*_*c*_ is the bacterial size. The effective viscosity of the liquid film is a strong function of the volume fraction of bacteria. The Krieger–Dougherty type law for effective viscosity is η = η_0_ (1-φ/0.466)^-2.6^, where φ is the volume fraction and η_0_= 10^−3^ Pa.s. The equation gives very close values to the effective viscosity calculated using the above-given formula for dewetting speed^48^. For the wetting velocity of order 10^−5^ m/s and 10^−6^ m/s, the effective viscosity of the medium is 0.0236 Pa.s. and 0.236 Pa.s, respectively. The corresponding bacterial volume fraction during dewetting is between 0.33 and 0.4. However, for bacteria, the curve fitting coefficients for calculating effective viscosity would vary, and there are no studies on the effective viscosity of bacterial solutions at volume fractions more than 0.1.

As shown in Fig. 5 (c), the SEM images are used to analyze the bacterial size and packing. The AFM image shows that the gap between the bacteria in the central regions is more than that at the edge. As a result, the Voronoi area of the bacteria aggregates in the central regions is higher than the edge bacteria aggregates, as shown in Fig. 5 (e). The variation in aspect ratio is nearly the same for the central and the edge regions. Since the bacteria in the central regions are also getting squeezed during the hole merging. However, the gap between the bacteria is the cause of a higher level of desiccation. The bacteria at the rim of the cellular deposits are also folded like the ones in the inner edges shown in Fig. 5 (d).

### D. Mechanistic insights

Along with the dewetting forces, the capillary forces due to the decaying liquid meniscus between bacteria make the bacteria move closer. Capillary forces led colloidal particle structuring in thin films is studied by Denkov and Kralchevsky et al.^45,49^. When the liquid film thickness decreases below the bacterial length scale, the drag force due to liquid flow decreases; microscopic video (Video 2) shows minimal bacterial movement. During the cellular pattern formation, the movement of the bacteria is driven by the capillary forces due to varying meniscus across the bacteria. In partially immersed particles, the deformation of the liquid meniscus gives rise to interparticle attraction or repulsion. The shape of the liquid meniscus between the bacteria and the surrounding region determines if the bacteria come closer or are moving apart. Capillary forces arise due to the pressure and surface effects at the three-phase contact line inclined at the particle surface. The capillary force of attraction is given by F_r_ = 2π σ r_c_^2^ (sin^2^φ_c_) (1/L) for r_c_ << L << (σ/ (Δρ)g)^1/2^, where r_c_ is the radius of the three-phase contact line at the bacterial surface, φ_c_ is mean meniscus slope angle at the contact line, σ is the surface tension of the liquid film and α is the contact angle of the bacteria. The distance L between the bacteria can be determined by the Laplace equation of capillarity as L is the length scale of bacteria.. The capillary force between the bacteria is of the order 10^−4^ N to 10^−6^ N, depending on the distance between the bacteria.

In the central regions, the bacteria do not initially settle down on the substrate because both the substrate and bacteria are negatively charged (electrostatic repulsion). So, the monolayer of bacteria would start to form when the thickness of the liquid layer is equal to the size of the bacteria. The immersion capillary forces between partially immersed bacteria are much higher than the floatation capillary forces. Thus, the varying meniscus across the bacteria in the film results in a downward capillary force on the bacteria. At a later stage, the evaporating liquid between the bacteria alters the liquid meniscus creating an attractive force bringing the bacteria together. For the case studied, only a monolayer of deposits has formed. At a higher bacterial concentration and slower evaporation rate, bilayer and trilayer deposits could form.

Fig. S4(a) shows a monolayer deposit of bacteria in the central regions. Malla et al.^50^ studied the effect of particle concentration and particle size on the edge deposition profile. The edge deposition is either a discontinuous or a continuous monolayer depending on the size of the particle, for very low particle concentrations below 0.1% v/v and diameter above 1 μm. For very low bacterial concentrations of less than 0.1% v/v and an effective diameter of more than 1 μm used in this study it is reasonable to obtain a monolayer deposition at the edge. However, the variation in the thickness (Fig. 4(b)) of the deposition can be attributed to varying levels of desiccation within the edge deposition and the squeezing.

### E. Bacterial survivability and pathogenesis aftermath desiccation of the droplet

We studied the survivability and infectivity of bacteria suspended in both Milli-Q water and surrogate respiratory fluid (SRF). Milli-Q water is used to have a clear visualization of bacteria which is inhibited in the presence of other colloids such as mucin in real biofluids. To compare the viability of bacteria in different regions, we scrapped the deposits at the edge keeping only the central deposits and vice versa. The dried droplet deposits, either whole droplet (W), edge removed (ER), or center removed (CR), are used for the viability and infectivity studies(refer to Fig.S5). We observed that the bacterial survival KP was higher in the edge (of the droplet) deposited bacteria compared to center deposited bacteria, both in the case of bacteria resuspended in Milli-Q water or SRF. We also observed that the infectivity of the bacterium deposited at the edge is significantly higher than the center deposited bacteria. This could be primarily due to the upregulation of virulence genes upon enormous force exerted on the bacteria during the drying process ^51^. We observe these similar observations for both SRF dried droplet and milliQ dried droplet, suggesting a possible role of physical force on the heightened virulence of the bacterium.

### F. Conclusions

The paper compares the nature of bacterial self-assembly, morphology, and viability in drying droplets. The bacterial cells generally die by desiccation. Studies^29,31^ have reported that desiccation is an important mechanism by which the bacterial cells in the drying sessile droplet die. This study shows that the self-assembly and desiccation of bacteria suspended in a drying droplet significantly alter the morphology of the bacteria. The bacteria packed closely experience lesser desiccation as their exposed area to the ambient is less. The above discussion serves as the reason why the bacterium at the edge is more viable than that at the central regions. In real biofluids and saline drying droplets, the increase in salt concentration causes osmotic shock, driving out water from the bacteria and resulting in death. In contrast, in a hypotonic solution like pure water used in this study, lysis could cause death. (by water intake and bursting). If bacteria survive the lytic process, water intake will help sustain the desiccation process longer than normal bacteria. A typical cellular pattern is formed by bacterial aggregation in the central region. The thin film instability led to the rupture of the liquid film causes, hole formation, and growth driving bacteria to come together and form cellular patterns. However, the gap between the bacteria is more in the central region, and the exposed area for desiccation is also higher. Thus, lesser viable bacteria are present in the central regions. Viability studies on realistic biofluids like respiratory fluid droplets also show a similar trend. A detailed study using actual biofluids is required to understand the phenomena better for practical applications.

## Methods

### Bacterial culture and cell line maintenance

The bacterial culture of *Klebsiella pneumoniae* (KP) was cultured in Luria Bertani (LB) broth overnight at 175 rpm shaker and 37°C incubators. The overnight grown (stationary phase) bacteria were washed five times with Milli-Q water to remove any remnants of the LB media and then resuspended in either Milli-Q water or SRF as mentioned in the particular experiment. For the Log phase, bacterial culture, the bacteria were sub-cultured at 1:33 ratio of bacteria: fresh media and incubated for 2 hours further at 37°C incubator shakers. For staining with a membrane dye called FM 4-64, the bacteria from stationary or log phase were incubated at the ice with dye in milliQ (at a concentration 1μg/mL of FM 4-64) for about 15mins, and bacterial cells were further washed with Milli-Q to remove excess dye and resuspended in Milli-Q or SRF as mentioned in a given experiment.

Murine macrophages were cultured in Dulbecco’s Modified Eagle’s Media (DMEM) containing 10% FBS at 37 °C incubator with 5% CO_2_. 12 hours prior to infection experiments, cells were seeded in 24 well plate at a confluency of 70-80%.

### Bacterial Viability and Gentamicin Protection Assay

The dried droplet, either whole droplet (W), edge removed (ER), or center removed (CR), was resuspended in PBS, and then the resuspended bacteria were plated onto LB agar to an enumerated viable bacterial number and was also used to infect in RAW264.7 macrophages, the cells were then centrifuged at 700rpm to increase the adherence of bacteria to mammalian cells. Next, the bacteria were incubated at 37°C incubators with 5% CO2. The cells were washed after 30mins with PBS, and media was removed and fresh media added containing 100ug/mL of gentamicin; further, after 1hour, media was removed and fresh media containing 25ug/mL of gentamicin. At 2hours and 16 hours post-infection, the mammalian cells were lysed and plated onto an LB agar plate to enumerate bacterial numbers at specific timepoints.

### Droplet Evaporation

The Surrogate respiratory fluid is prepared according to the procedure mentioned in Rasheed et.al.^52^The bacteria suspended in Milli-Q and SRF droplets are casted on the glass substrate cleaned with Isopropyl alcohol. During the experiments, ambient conditions are maintained at the relative humidity 50 ± 3% and temperature 27 ± 1^°C^. Droplets of volume 0.95 ± 0.1 μl are drop cast on glass substrates cleaned with Isopropyl alcohol. The droplets showed an initial contact angle of 46 ± 2^°^; the droplet dries in pinned contact line mode with a wetting diameter of 2.2 ± 0.07 mm. The bacterial motion and deposition near the end of evaporation are recorded using the bright and dark field optical method.

A bright-field image of the final deposition pattern is recorded using a DSLR camera. Scanning electron microscopy imaging reveals the finer details of the deposition and the arrangement of bacteria.

### Cell imaging and live tracking

The bacterial motion is tracked using membrane stain FM 4-63 with a excitation/emission maximum of 515/640 nm at 2.28 fps using a Leica TCS SP8 microscope.

### Micro-Particle Image Velocimetry (μ-PIV)

The μ-PIV technique is used to qualitatively and quantitively study the flow inside droplets. Neutrally buoyant monodisperse (of size 860 ± 5 *nm*) fluorescent polystyrene particles are used for *μ*-PIV experiments. The polystyrene particles mentioned above were procured from ThermoFisher Scientific. These fluorescent polystyrene particles are added to the bacteria-containing liquid at 0.008% by volume. These polystyrene particles faithfully follow the flow as their stokes number is (St) 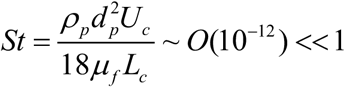. Where, *ρ*_*p*_ is the particle density, *d*_*p*_ is the diameter of the particle, *µ*_*f*_ is the fluid viscosity, *L*_*c*_ is the characteristic length scale (∼contact diameter of the droplet), *U*_*c*_ is the expected velocity scale. The droplet containing bacteria (non-fluorescent) and polystyrene particles (fluorescent) is volumetrically illuminated using an Nd: Yag laser (NanoPIV, Litron Laser). Images are captured from the bottom of the solvent droplet using an Imager Intense camera fitted to a Flowmaster MITAS microscope [Field of View: 1000 × 600 *μm*^2^, depth of field: 28 *μm*]. The images are acquired at 1 *fps* using a single frame–single pulse technique. The particles move 3-4 pixels in the subsequent frames, optimal for PIV computation. All images are pre-processed by applying appropriate background subtraction. Post-processing is done by pair-wise cross-correlation of sequential single-frame images. The interrogation window is maintained at 64 × 64 pixels for the first pass and 32 × 32 pixels for the subsequent passes with 50% overlap between two windows. The instantaneous vector fields thus obtained are temporally averaged (over the time period pertaining to each regime) to obtain the final velocity vectors. The above post-processing of images is done using DaVis7.2 software from LaVision GmbH.

### Atomic force Microscopy

Park NX10 atomic force microscope is used to obtain the images and force distance spectroscopy analysis. The CONTSCR probe with 0.2 N/m stiffness is used for the measurements. The nanoindentation is done at the speed of 0.3 μm/s for both up and down directions without holding.

### Credit statement

Conceptualization: S.B. and D.C.; Methodology: S.B., D.C., A.R., O.H., R.C., S.R.S.; Investigation: A.R., O.H., R.C., S.R.S.; Visualization: A.R., O.H., R.C., S.R.S.; Funding acquisition: D.C. and S.B.; Project administration: D.C. and S.B.; Supervision: D.C. and S.B.; Writing— original draft: A.R., O.H., R.C., S.R.S.; Writing: A.R., O.H., R.C., S.R.S.; editing and revision: S.B., D.C., A.R., O.H., R.C., S.R.S.

### Data availability statement

All relevant data are within the paper and its Supporting Information files. All materials and additional data are available from the corresponding author upon request.

