## Supplementary Material for "Physics of Self-Assembly and Morpho-Topological Changes of *Klebsiella Pneumoniae* in Desiccating Sessile Droplets"

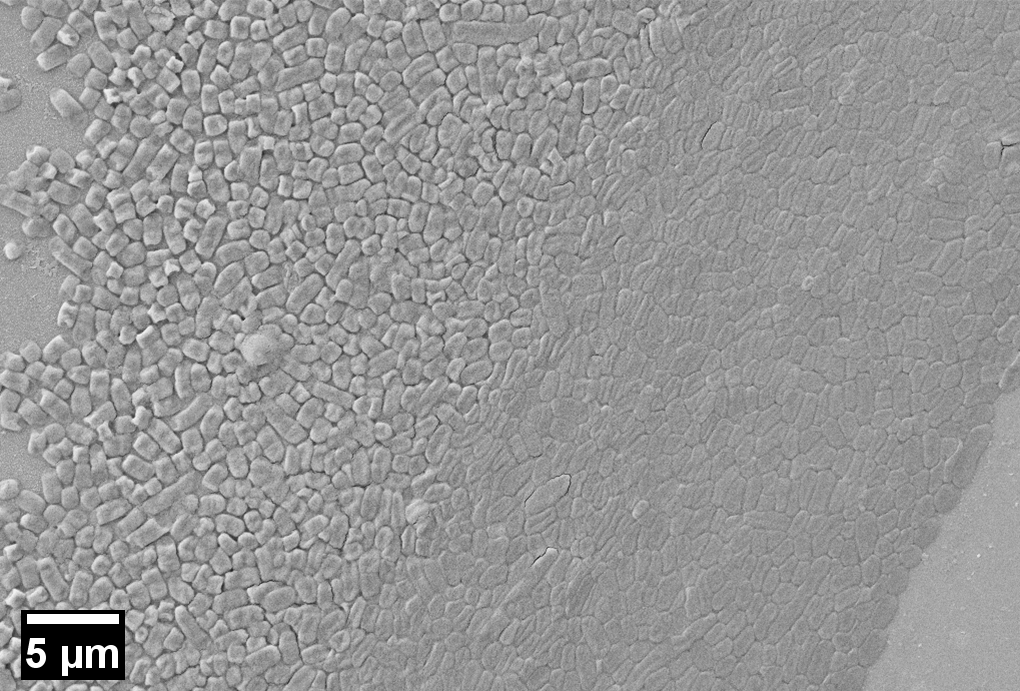
**(a)**

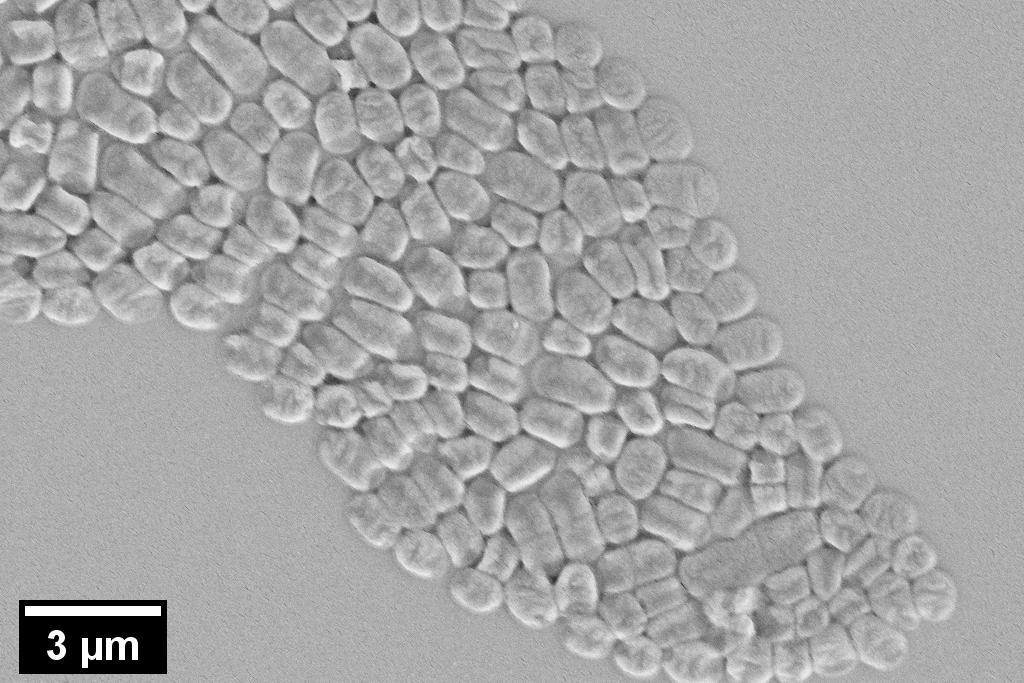
**(b)**

**Fig.S1: Scanning electron microscopy of the bacterial deposits (a) at the edge and (b) at the central region**

**Table 1: Zeta potential measurement of Klebsiella pneumonia**

| **Run** | **Mobility** | **Zeta potential (mV)** | **Relative Residual** |
| --- | --- | --- | --- |
| 1 | -2.51 | -32.12 | 0.0306 |
| 2 | -2.07 | -26.46 | 0.0314 |
| 3 | -2.31 | -29.51 | 0.0222 |
| 4 | -2.56 | -32.76 | 0.0171 |
| 5 | -2.37 | -30.3 | 0.0156 |
| **Mean** | **-2.36** | **-30.23** | **0.0234** |
| **Std.Error** | 0.09 | 1.11 | 0.0033 |
| **Combined** | -2.36 | -30.14 | 0.0074 |

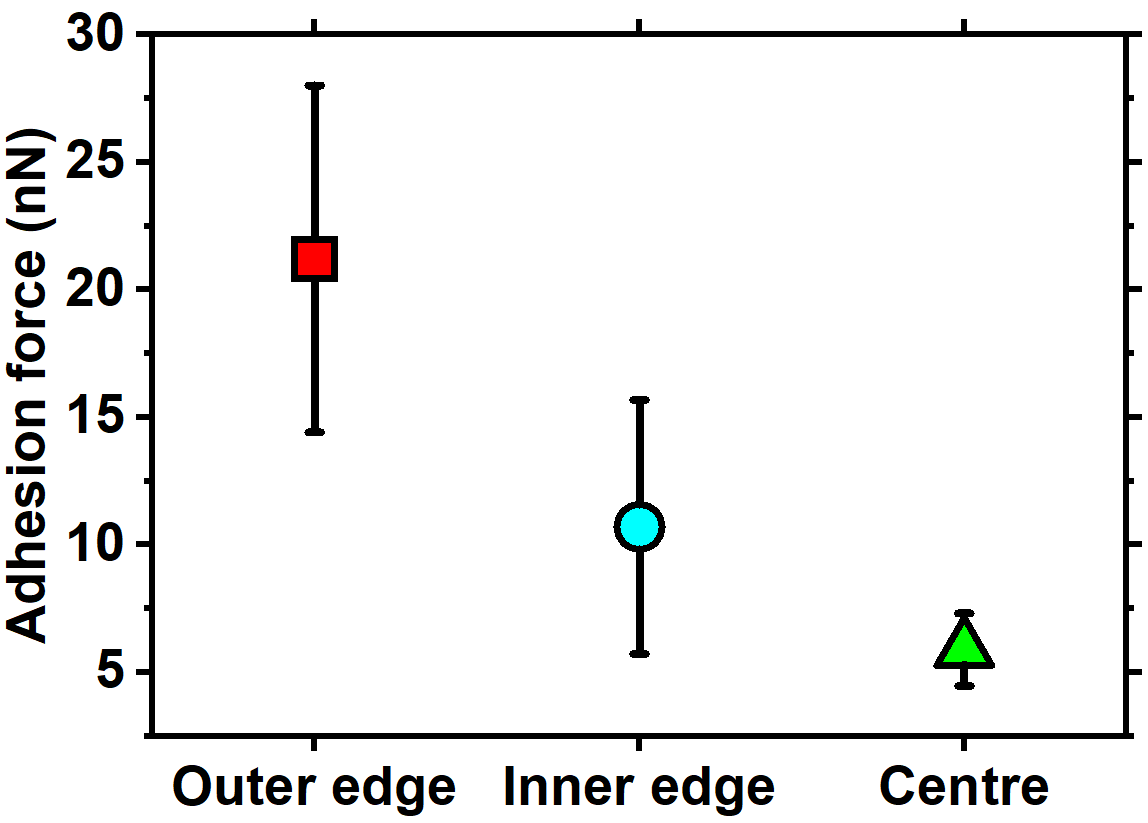

**Fig.S2: Adhesion force of bacterial deposits at different locations.**

**Analysis of hole growth during cellular pattern formation**

Only single unmerging holes are taken into consideration, while few holes merge and become large holes. The red-coloured symbols correspond to the normalized area of different holes that grow adjacent to the coffee ring. The blue-coloured symbol corresponds to the holes in the central region. The different symbols correspond to different cases.

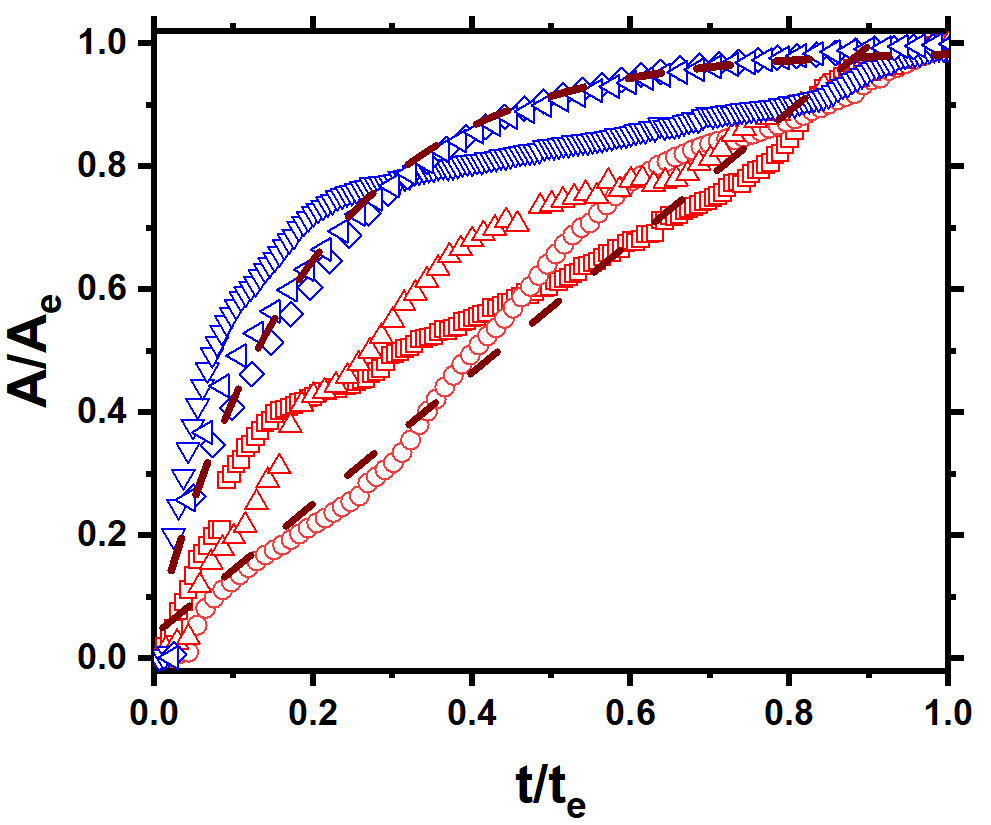

**Fig. S3: Normalized instantaneous hole area to the normalized time, t is the time since the inception of the hole, and t_e_ is the total time for a hole to grow to its maximum.**

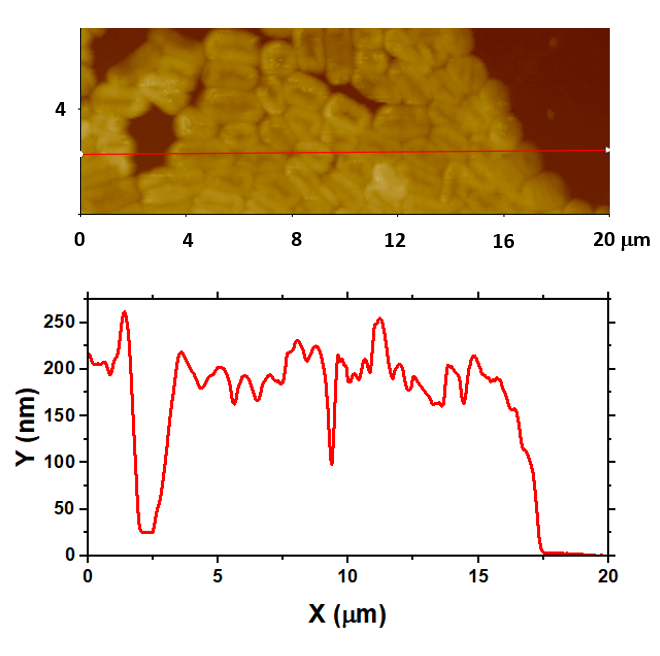

**Fig.S4 (a): Thickness profile at the central region.**

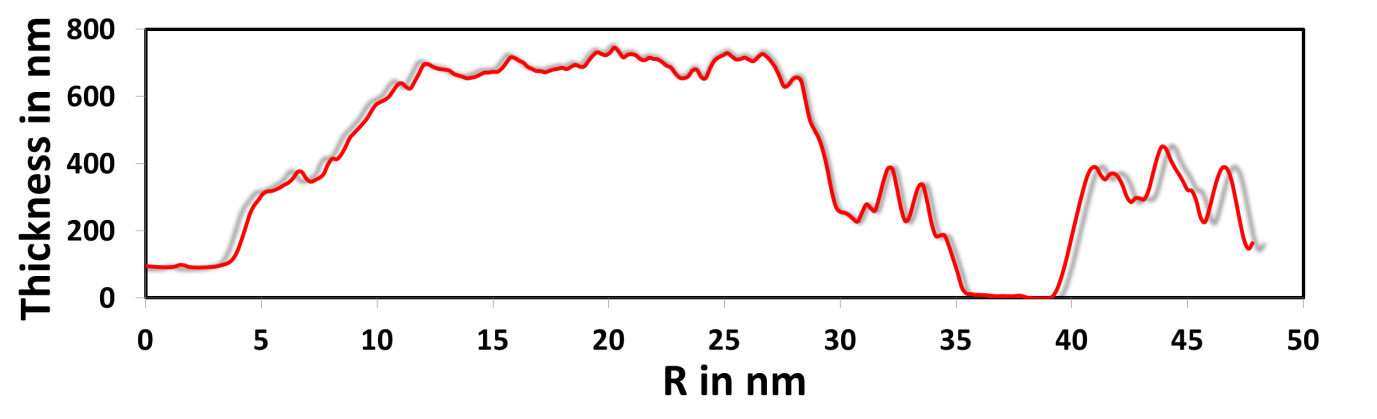

**Fig.S4 (b): Thickness profile of the edge deposition vs R, the distance from the outermost edge.**

**Viability and infectivity analysis of central and edge deposits**

The deposits are resuspended according to the procedure mentioned in the methods section. We compared the viability of the bacterial deposits at edge and the central deposits with respect to the whole deposits by removing the central and edge deposits respectively. The deposits are scrapped off using a fine injection needle. The corresponding deposits left off by removing the edge and the central regions is shown in Fig.S5. twenty-seven such deposits per case per culture is considered for the viability and infectivity tests.

**Centre removed, CR**

**Edge removed, ER**

**Scrapping**

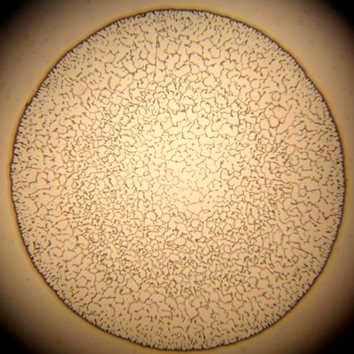

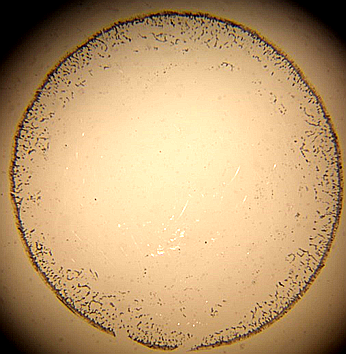

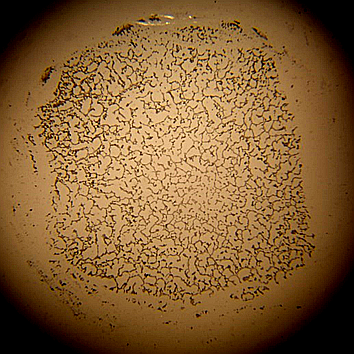

**Fig. S5: Bacterial deposits scrapped for viability test.**
